# Deep learning-based image classification reveals heterogeneous execution of cell death fates during viral infection

**DOI:** 10.1101/2024.10.03.616527

**Authors:** Edoardo Centofanti, Alon Oyler-Yaniv, Jennifer Oyler-Yaniv

## Abstract

Cell fate decisions, such as proliferation, differentiation, and death, are driven by complex molecular interactions and signaling cascades. While significant progress has been made in understanding the molecular determinants of these processes, historically, cell fate transitions were identified through light microscopy that focused on changes in cell morphology and function. Modern techniques have shifted towards probing molecular effectors to quantify these transitions, offering more precise quantification and mechanistic understanding. However, challenges remain in cases where the molecular signals are ambiguous, complicating the assignment of cell fate. During viral infection, programmed cell death (PCD) pathways, including apoptosis, necroptosis, and pyroptosis, exhibit complex signaling and molecular crosstalk. This can lead to simultaneous activation of multiple PCD pathways, which confounds assignment of cell fate based on molecular information alone. To address this challenge, we employed deep learning-based image classification of dying cells to analyze PCD in single Herpes Simplex Virus-1 (HSV-1)-infected cells. Our approach reveals that despite heterogeneous activation of signaling, individual cells adopt predominantly prototypical death morphologies. Nevertheless, PCD is executed heterogeneously within a uniform population of virus-infected cells and varies over time. These findings demonstrate that image-based phenotyping can provide valuable insights into cell fate decisions, complementing molecular assays.

## Introduction

Cell fate decisions—such as proliferation, differentiation, or death—are the result of complex molecular interaction networks and signaling cascades. Although today we know a great deal about the molecular species that signify major cell fate and state transitions, historically, such transitions were visualized by microscopy and defined by changes in morphology or function.

As our understanding of the signaling pathways governing cellular decisions deepened, morphology-based classification gradually gave way to techniques that probe for key molecular effectors. These assays offer precise, quantifiable biological data, whereas cell images remain complex, heterogeneous, and often require specialized expertise to interpret. However, despite the advantages of molecular probe-based assays, signaling can be ambiguous and context-dependent, making it difficult to assign cell phenotype or fate from molecular data alone.

Ambiguity in signaling can arise from various factors: technical limitations, where some molecules are challenging to measure accurately; biological promiscuity, where similar downstream signals can be activated by different upstream triggers; biological stochasticity, where fate commitment is driven by inherently unpredictable biochemical noise; or contextual variations, where the same signals may lead to different outcomes in distinct environments. Furthermore, many cell fate decisions are driven by multiple signals that are difficult to measure simultaneously in single cells.

We previously studied the timing of commitment to programmed cell death (PCD) during viral infection, and discovered that it is well-described by a memoryless exponential process (Oyler-Yaniv *et al*., 2021). Consequently, the fates of individual cells cannot be deduced from their current state. PCD is a conserved form of defense against viruses and other intracellular pathogens, which effectively short circuits the pathogen’s life cycle and restricts spread (Jorgensen *et al*., 2017). Aside from the timing of cell death, the PCD decision involves a choice between multiple different PCD types.

The three major types of PCD that occur during viral infection - apoptosis, necroptosis, and pyroptosis - differ in the molecular circuits that control their execution, cell morphology preceding death, and on their ability to stimulate inflammation (Ketelut-Carneiro and Fitzgerald, 2022). While apoptosis is considered anti-inflammatory or immunologically silent, necroptosis and pyroptosis, both lytic forms of PCD, are highly inflammatory due to the release of proinflammatory cytokines and danger-associated molecular patterns (DAMPs). Elevated levels of lytic cell death during infection are associated with more severe symptoms, impaired tissue function, and hyper-inflammation (Rodrigue-Gervais *et al*., 2014; Xu *et al*., 2017; Karki *et al*., 2021, 2022; Junqueira *et al*., 2022; Sefik *et al*., 2022; Liang *et al*., 2024). It is therefore important to be able to measure – and potentially control – the execution of PCD fates during viral infection. However, there is significant crosstalk and promiscuity in PCD signaling, making it challenging to ascertain how cells ultimately die (Snyder and Oberst, 2021).

Given that unique and stereotypic death-associated morphologies are defining characteristics of each PCD type, we reasoned that morphology may be useful to distinguish cell death modes during infection (Ketelut-Carneiro and Fitzgerald, 2022). However, extracting morphological information from images of cells in a systematic and scalable manner is difficult. In recent years, deep learning-based image classification tools have revolutionized computational image analysis (Moen *et al*., 2019). In instances where ground truth data are available, models can be trained to make classifications directly from image pixels. This presents an opportunity to automate—and therefore dramatically scale up—cellular phenotyping that is otherwise accomplished manually by trained personnel. These models offer rapid and accurate performance on routine tasks such as automated segmentation of nuclei and dead cell identification (La Greca *et al*., 2021; Stringer *et al*., 2021). More recently, they have been applied to identify more subtle and complex cellular phenomena such as cell cycle, senescence, stem cell differentiation, and metastatic potential (Waisman *et al*., 2019; Zaritsky *et al*., 2021; Heckenbach *et al*., 2022; Jose *et al*., 2024).

Motivated by these rapid advancements, we used a combination of microscopy and deep learning-assisted image analysis to study PCD signaling and execution in single Herpes Simplex Virus-1 (HSV-1)-infected fibroblasts. We find that virus-infected cells often activate multiple PCD signaling pathways simultaneously. Despite promiscuity in signaling, dying cells adopt heterogeneous, yet prototypical morphologies, indicating commitment to a particular death fate. Finally, we tested the hypothesis that the different PCD pathways are functionally redundant, such that cells can flexibly switch between them. However, blocking of caspases, which drive apoptosis, led to fractional killing, whereas blocking receptor-interacting protein kinases 1 (RIP1), which drives necroptosis, delayed execution of both apoptosis and necroptosis. In neither case did we observe functional compensation by the intact pathways. In summary, we used deep learning-based image classification of PCD fates during infection to overcome limitations imposed by ambiguity in the upstream signaling network. We expect this strategy to continue enabling new insights in cell fate decisions rooted in cell morphology, in addition to molecular machinery.

## Results

### Single, virus-infected cells activate more than one death signaling pathway simultaneously

Infection with Herpes Simplex Virus-1 (HSV-1) has been reported to activate both apoptosis and necroptosis depending on the cell type and context (Guo *et al*., 2015; Huang *et al*., 2015). Further, bulk pull-down assays have shown that HSV-1 and other pathogens can lead to the physical association of effector proteins from both death pathways (Lee *et al*., 2021). These observations suggest that during infection, both apoptosis and necroptosis may be activated simultaneously. We therefore wanted to examine the activation of apoptotic and necroptotic signaling pathways within single HSV-1-infected cells.

To do so, we visualized the activation of terminal effector proteins in the apoptotic and necroptotic signaling pathways: cleaved caspase 3 (Cl Casp 3) and phosphorylated mixed lineage kinase domain-like protein (pMLKL) (Galluzzi *et al*., 2018). During apoptosis, procaspase 3 is cleaved into its active form by the initiator caspase 8. Cl Casp 3 is referred to as the executioner caspase because it digests many cellular proteins during the final stage of apoptosis. During necroptosis, MLKL is phosphorylated by the receptor-interacting protein kinases 1 and 3 (RIP1/3), leading to its oligomerization and pore formation in the plasma membrane, ultimately resulting in cell lysis.

As controls, we visualized Cl Casp 3 and pMLKL in mouse 3T3 fibroblasts treated with tumor necrosis factor α (TNFα), an initiator of extrinsic apoptosis; doxorubicin, an activator of intrinsic apoptosis; or TNFα combined with the pan-caspase inhibitor Z-Vad(Ome)-FMK (ZVad), which activates necroptosis (Vercammen *et al*., 1998; Gewirtz, 1999; Li and Beg, 2000; Micheau and Tschopp, 2003; Zhang *et al*., 2009). Our data show that each treatment causes activation of either Casp 3 or MLKL, but not simultaneous activation of both (Fig 1A).

**Figure 1.**
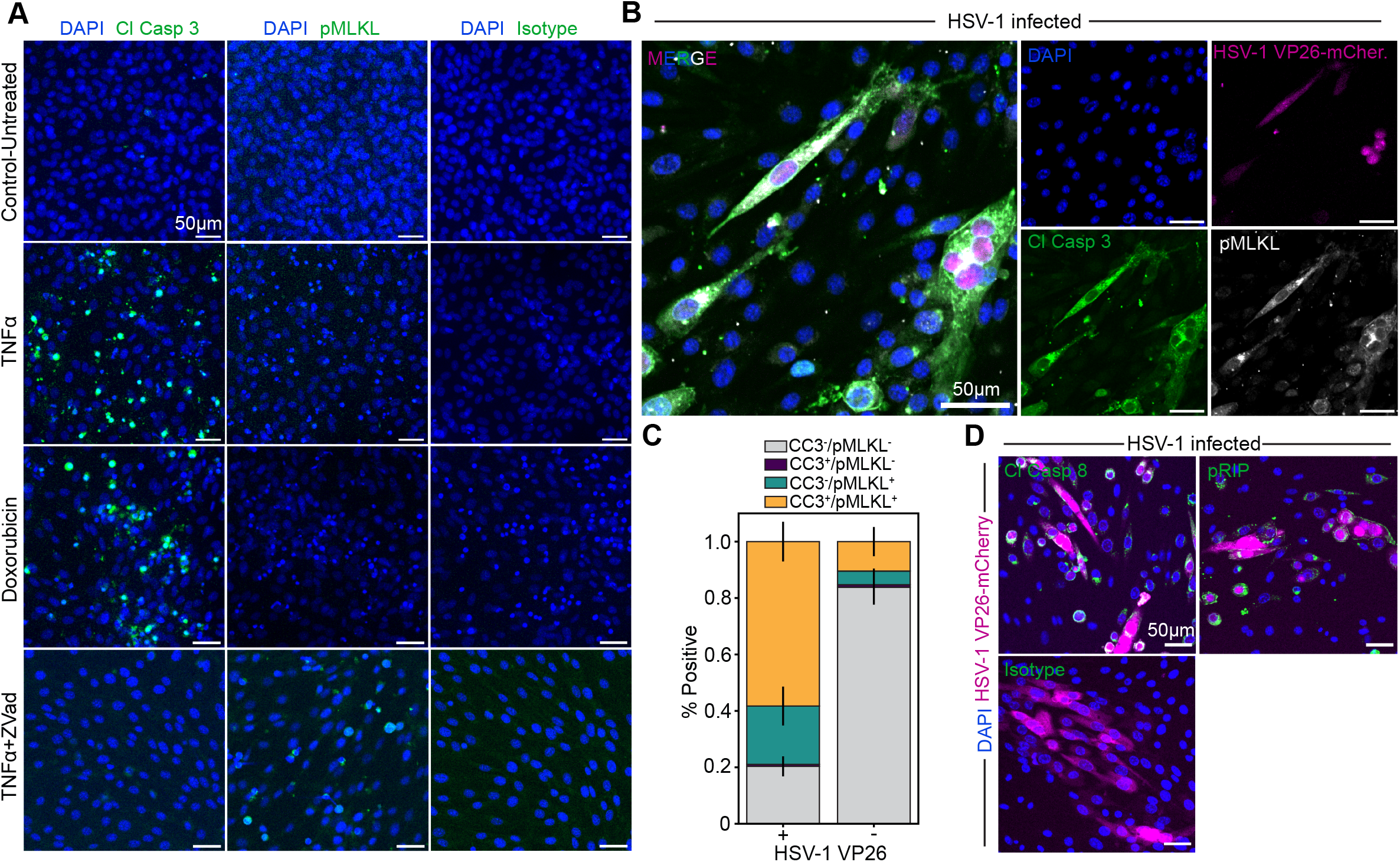
Virus-infected cells simultaneously activate both apoptotic and necroptotic signaling pathways. **A**. Representative immunofluorescence images of mouse fibroblasts treated with the indicated compounds and stained for either cleaved caspase 3 (Cl Casp 3), phosphorylated MLKL (pMLKL), or an isotype control. **B**. Representative immunofluorescence images of HSV-1 infected mouse fibroblasts stained for both Cl Casp 3 and pMLKL simultaneously. **C**. Quantification of the total proportion of either HSV-1 VP26-positive or negative cells that are positive for either PCD signaling marker, or both markers. The error bars show the standard error of the mean of four independent experiments. **D**. Representative immunofluorescence images of HSV-1 infected mouse fibroblasts stained for either cleaved caspase 8 (Cl Casp 8), or phosphorylated RIP1 (pRIP). All images are representative of results from at least three independent experiments.

We next infected fibroblasts with a multiplicity of infection (MOI) of 2 with Herpes Simplex Virus-1 (HSV-1), strain KOS-VP26-mCherry, where mCherry is fused to the viral small capsid protein VP26 (Etienne *et al*., 2017). Immunofluorescence staining revealed that, unlike the control treatments, a majority of VP26-positive infected cells stain positive for both Cl Casp 3 and pMLKL (Fig 1B-C). Eighty-five percent of VP26-negative cells are negative for all programmed death pathways; however, a small percentage are positive for both markers or pMLKL alone. As VP26 is a late-stage viral protein, it is likely that some VP26-negative cells that stain positive for death markers are in early stages of infection that cannot be visualized using VP26 reporter viruses (Knipe and Howley, 2007). This is supported by the observation that uninfected control cells are negative for death markers, and that a fraction of cells negative for viral VP26-mCherry stain positive for the immediate-early viral protein ICP4 (Fig 1A, Fig S1A-B). Staining for upstream markers of the apoptotic and necroptotic pathways—Cl Casp 8 and phosphorylated RIP1 (pRIP), respectively—further supports the idea that these pathways are heterogeneously activated, both across a population of virus-infected cells and within individual infected cells (Fig 1D).

Taken together, these data demonstrate that a significant proportion of individual HSV-1-infected cells simultaneously activate both the apoptotic and necroptotic signaling pathways. This overlap in cell death signaling makes it difficult to determine how infected cells ultimately execute cell death. Given that different forms of PCD are associated with unique and distinctive morphologies, we asked whether cell death-associated morphologies could be used to determine the mode of death.

### Pharmacologic induction of each of the three major PCD types

To study the execution of PCD during viral infection, where heterogeneity is anticipated, we first established a system to observe the morphologies of each death type in isolation. Although only apoptosis and necroptosis are associated with HSV-1 infection in non-immune cell types, we also included pyroptosis-inducing compounds in our studies, since published transcriptomic and proteomic datasets confirm that our cells express the core components of all three pathways (Fig S2A-B) (Schwanhäusser *et al*., 2011).

Next, we treated cells with established pharmacologic treatments that result in uniform activation of each death type (referred to as ‘prototype conditions’). To induce apoptosis, we treated cells with Tumor Necrosis Factor-α (TNFα), the translation inhibitor cycloheximide (CHX), and a specific inhibitor of RIP1K, Necrostatin-1 (Nec1) (Fig S2C) (Degterev *et al*., 2005, Degterev 2008). To induce necroptosis, we treated cells with TNFα, CHX, and the caspase inhibitor ZVad (Vercammen *et al*., 1998; Li and Beg, 2000; Zhang *et al*., 2009). In both treatments, CHX promotes the decay of the labile protein c-FLIP, which antagonizes the formation of both the death-inducing signaling complex and the ripoptosome (Irmler *et al*., 1997; Feoktistova *et al*., 2011; Roux *et al*., 2015). To activate pyroptosis, we treated cells with a combination of bacterial lipopolysaccharide (LPS) and nigericin, a potassium ionophore that leads to inflammasome activation (Mariathasan *et al*., 2006).

Each of the cytotoxic prototype treatments induces cell death, yet to varying extents and with varying kinetics (Fig 2A-B). While the initial onset of death is slightly earlier for necroptosis compared to apoptosis, both treatments result in comparable overall death by 48 hours, and the overall death rate is faster for apoptosis. By contrast, pyroptosis is slower, likely reflecting the time needed to upregulate some components of the pathway, accomplished via the LPS priming signal (Bauernfeind *et al*., 2009). Finally, we quantified extracellular ATP in response to the different cytotoxic treatments. ATP is a canonical DAMP released during lytic death that binds to purinergic receptors to mediate inflammation (Linden *et al*., 2019). This experiment demonstrates that both necroptosis- and pyroptosis-inducing treatments cause significantly greater ATP release than apoptosis, reflecting the lytic nature of these forms of cell death (Fig 2C). These data demonstrate that while 3T3 fibroblasts are predisposed to apoptosis, under different conditions, they are competent to undergo necroptosis and pyroptosis, which lead to the release of inflammatory DAMPs.

**Figure 2.**
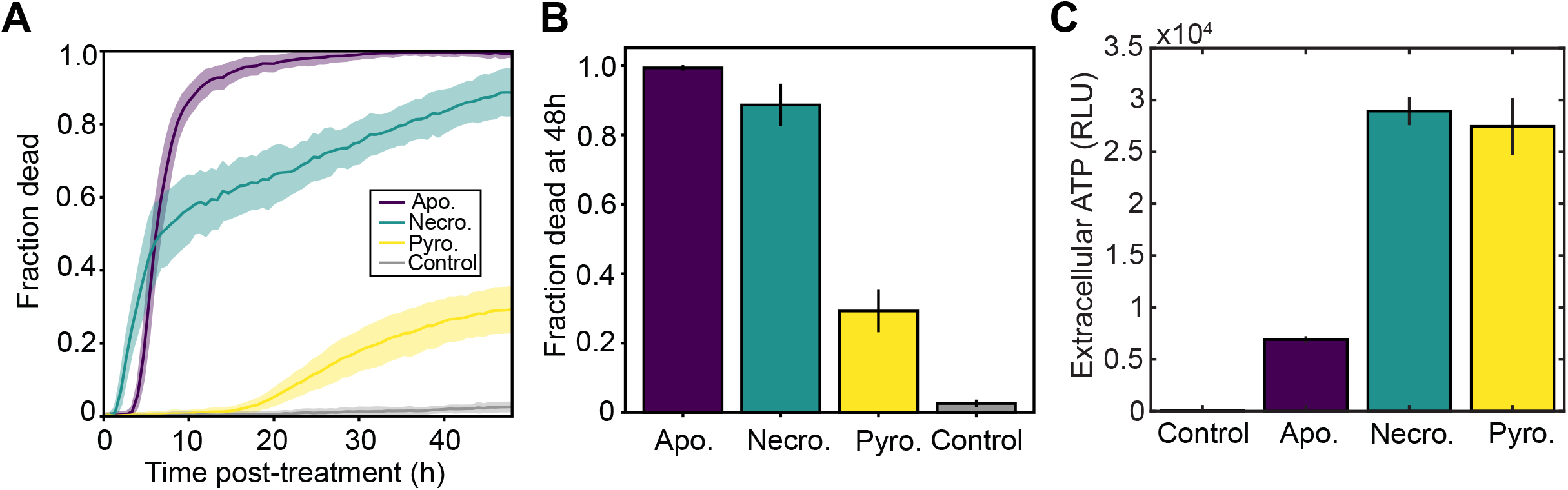
Pharmacologic induction of each PCD type. **A**. Quantification of cell death over time for each of the different prototype PCD inducing treatments (Apo: TNFα, CHX, Nec1; Necro: TNFα, CHX, ZVad; Pyro: LPS and Nigericin). **B**. Quantification of the total fraction of cells dead at 48 hours post-treatment. **C**. Quantification of extracellular ATP for each of the different prototype treatment conditions. Error bars show the standard deviation of multiple

### An image classifier distinguishes the different forms of PCD based on cellular morphology alone

The different modes of PCD are associated with distinct morphological characteristics (Ketelut-Carneiro and Fitzgerald, 2022). Apoptosis is characterized by membrane blebbing, nuclear shrinkage, and fragmentation, followed by the release of membrane-bound apoptotic bodies. Necroptosis is marked by swollen organelles, rounding of the cell body, and an abrupt rupture of the plasma membrane. Pyroptosis involves intracellular vacuolization and a characteristic “parachute” morphology. Both necroptosis and pyroptosis lead to pore formation—although by distinct pore-forming proteins—which results in osmotic stress, cell swelling, and lysis. To visualize the morphological differences between these types of cell death, we treated cells with cytotoxic prototype compounds and imaged them using differential interference contrast (DIC) microscopy. Each treatment resulted in consistent and distinct morphologies at the time of cell death that were visually discernible (Fig 3A, Supplemental movies 1-3).

**Figure 3.**
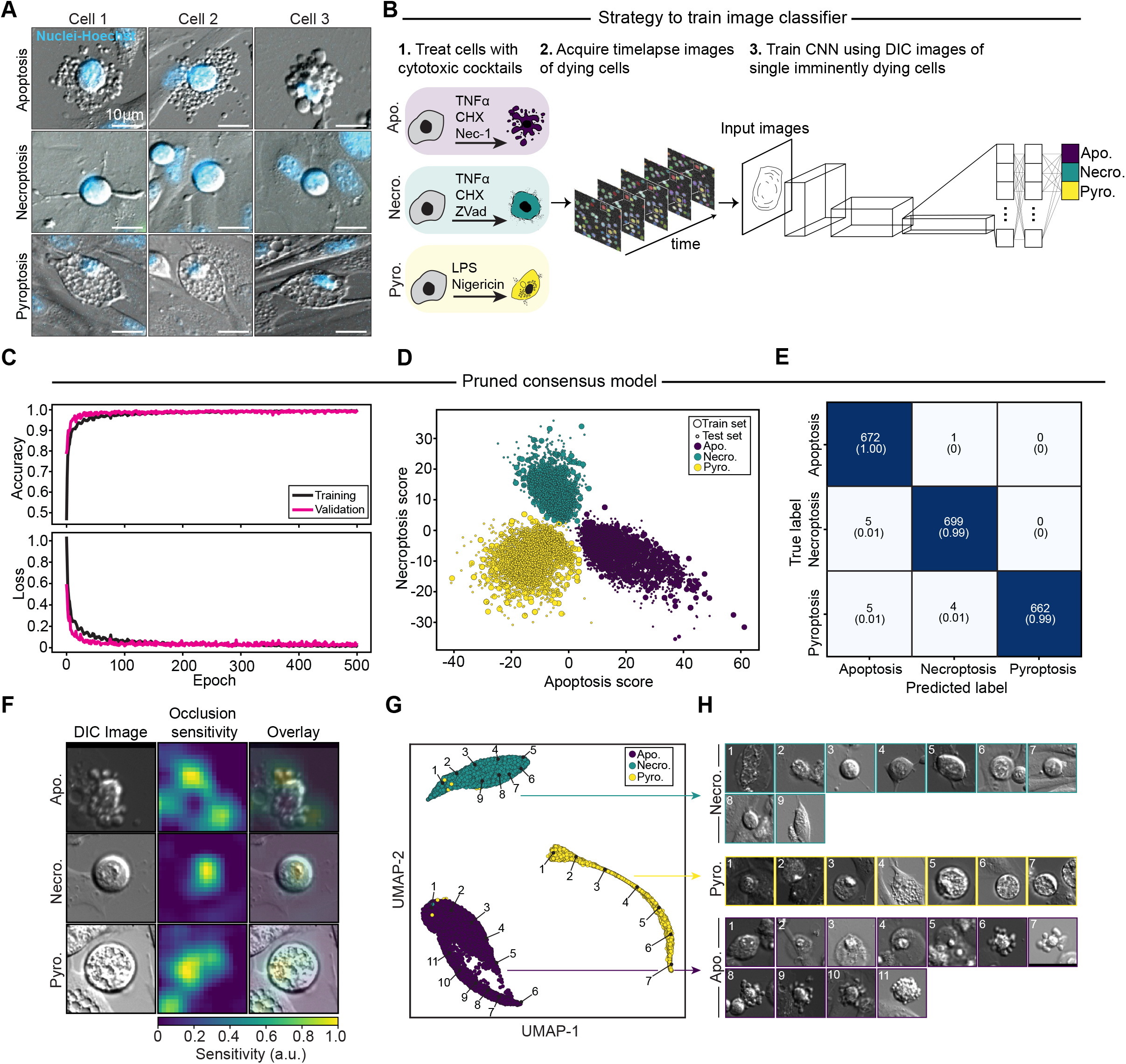
An CNN classifier distinguishes the different forms of PCD based on cellular morphology alone. **A**. Representative examples of imminently dying apoptotic, necroptotic, and pyroptotic cells from each of the different cytotoxic prototype conditions. **B**. Cartoon diagram showing the general approach for training a CNN-based image classifier. Cells are treated with compounds that uniformly induce each PCD type, then imaged over time. Then images of single, imminently-dying cells are used to train a CNN-based image classifier. **C**. Performance of the classifier during training following pruning of the training data via a multi-model consensus strategy. **D**. Logit scores of the train and test data following pruning. **E**. Confusion matrix demonstrating model performance following data pruning. Numbers represent the number of cells classified (top) and fraction called correctly (bottom). **F**. Occlusion sensitivity for representative cells from each death prototype class. **G**. UMAP dimensional reduction of feature space for cells from each class. **H**. Representative images of single cells sampled from the indicated points within the UMAP clusters. For all of figure 3, images of cells used to train, validate, and test the model were aggregated from three independent imaging experiments, each with at least 5 technical replicates per prototype condition.

Motivated by the observation of distinct death morphotypes, we trained the residual convolutional neural network (CNN) ResNet50 using images of cells that were imminently dying (Fig 3B) (He *et al*., 2015). Cells were first treated with cytotoxic prototype compounds and imaged over time. Dead cells were then identified based on a combination of either or both of the following: (1) an abrupt and sustained increase in nuclear Hoechst intensity, indicating nuclear condensation characteristic of apoptosis; and (2) incorporation of Sytox green, indicating loss of membrane integrity. An imminently dying cell was defined as the image of the cell from the frame just prior (within 40 m) to death. The images of imminently dying cells from each prototype condition were then used as labeled data to train our death mode classifier.

To enhance robustness, the training image datasets were augmented through cropping, rotations, reflections, and intensity adjustments before being applied to model training. After initial training, the datasets were pruned using a consensus strategy, where only images correctly classified by ten separately trained models were retained. This approach, a variation of Confident Learning, helps automatically identify and remove label errors in the dataset, ensuring the model is trained on accurate prototypical examples (Northcutt *et al*., 2019). The pruned dataset was then used to train a new model, which was subsequently validated and tested on unseen images of prototype cells. Pruning significantly improved both training speed and model performance, evidenced by clearer separation of logit scores between death prototypes and higher precision, recall, and accuracy (Fig 3C-E, Fig S3A-C, and Table I). Logit scores quantify the probability that a given cell falls into a particular death class. To determine which regions of the images were most critical for model classification, we measured the model’s sensitivity to occluding different image areas. Notably, the model was most sensitive to disruptions in the cell body, including the nucleus, cytoplasm, and apoptotic bodies, across all death prototype conditions (Fig 3F and Fig S3D-F). This indicates that the model used meaningful cellular morphology rather than irrelevant or spurious features for classification.

**Table I.** Model performance statistics.

| Table I. Model performance statistics |  |  |  |  |  |  |  |  |
| --- | --- | --- | --- | --- | --- | --- | --- | --- |
|  | Precision |  | Recall |  | F1 Score |  | Support |  |
| Apoptosis | 0.88 | 0.98 | 0.92 | 1.0 | 0.9 | 0.99 | 954 | 675 |
| Necroptosis | 0.88 | 0.99 | 0.88 | 0.99 | 0.88 | 0.99 | 1036 | 672 |
| Pyroptosis | 0.97 | 1.0 | 0.92 | 0.99 | 0.94 | 0.99 | 960 | 701 |
| Overall accuracy | 0.91 |  |  |  | 0.99 |  |  |  |
| Single model/model trained on consensus-pruned data |  |  |  |  |  |  |  |  |

We examined the diversity of death-associated morphologies within each prototype class. To do this, we analyzed the final fully connected layer of the trained model that contained 2048 learned features, and performed dimensional reduction (Uniform Manifold Approximation and Reduction - UMAP) on this feature space. This analysis demonstrates that cells within each prototype death class cluster separately from one another (Fig 3G). Examination of representative cells from within each cluster reveals in-class morphological diversity (Fig 3H). This diversity may arise from differences in the timing of cell death between frames, biological variability in the morphology of each PCD type, or a combination of both. In either case, our model correctly classifies cells dying via each prototypical death mode, even in situations where a human would struggle to do so. In summary, we developed a method for training a highly accurate CNN-based classifier capable of assigning death fate from microscopy images based on cellular morphology alone. Our trained classifier enables us to study variability in death-associated morphologies during viral infection, where heterogeneous death fates are expected.

### Dying, virus-infected cells adopt diverse and variable death-associated morphologies

Our previous work demonstrates that treatment of HSV-1-infected fibroblasts with a combination of Nec1 (RIP1 inhibitor) and ZVad (pan-Caspase inhibitor) dramatically reduces cytotoxicity (Oyler-Yaniv *et al*., 2021; Sonnett *et al*., 2024). This indicates that PCD, rather than unregulated necrosis, constitutes the majority of cell death during HSV-1 infection. To examine the morphology of dying, infected cells, we infected cells with HSV-1 strain KOS-VP26-mCherry at a multiplicity of infection (MOI) of 10 and imaged them over a 48-hour period. This high MOI ensures that ∼90% of cells are infected synchronously from the initial viral exposure, minimizing variability in the timing of infection (Oyler-Yaniv *et al*., 2021; Sonnett *et al*., 2024). Visual examination of death morphologies in infected cells reveals greater heterogeneity than that observed under prototype conditions (Supplemental movie 4). This indicates that cell death during viral infection is morphologically diverse. We next used our trained cell death image classifier to systematically examine this variability.

We projected the multi-dimensional feature space from images of dying cells onto a UMAP of prototype cells, color-coding based on the timing of death post-infection (Fig 4A). Most dying cells are morphologically similar to necroptotic and apoptotic prototype cells, rather than pyroptotic prototypes. This is consistent with the observation that pyroptosis is more common in innate immune cell types, such as macrophages, and that our fibroblasts are not predisposed to undergo pyroptosis (Fig 2A-B). Necroptosis is the predominant form of death early during infection, whereas apoptosis occurs later (Fig 4A). This is consistent with reports that ICP6, an early-expressed HSV-1 protein, directly interacts with mouse RIP1 and RIP3 to initiate necroptotic signaling (Huang *et al*., 2015). Visual examination of representative dying, HSV-1-infected cells from each cluster shows that cells are morphologically similar to dying prototype cells, and that there is comparable heterogeneity within each death morphotype (Fig 4B).

**Figure 4.**
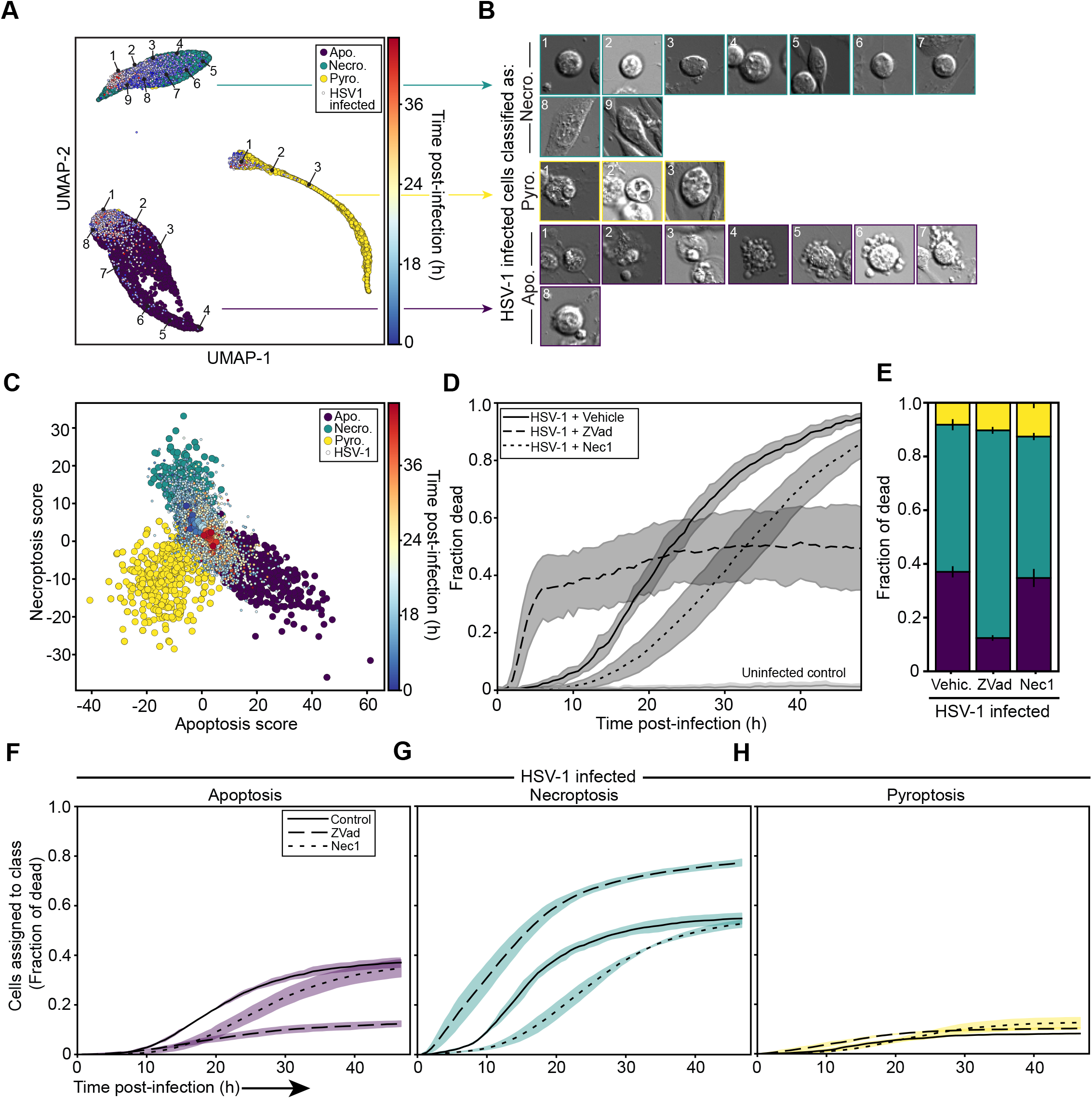
Virus-infected cells execute heterogeneous forms of PCD. **A**. UMAP dimensional reduction of feature space for cytotoxic prototype-treated cells, with virus-infected cells overlaid on top. **B**. Representative images of single cells sampled from the indicated points within the UMAP clusters. **C**. Logit scores of test data from cytotoxic prototype-treated cells, with virus-infected cells overlaid on top. Small time-colored circles represent single virus-infected cells, while the larger time-colored circles represent the average classification for each time point. **D**. Kinetics of cell death for virus-infected cells treated with the vehicle control, ZVad, or Nec1. **E**. Total proportion of virus-infected cells classified as apoptotic, necroptotic, or pyroptotic with or without ZVad or Nec1 treatment, over the course of the 48 hour experiment. **F-H**. Kinetics of cell classification as apoptotic (F), necroptotic (G), or pyroptotic (H) for virus-infected cells treated with the vehicle control, ZVad, or Nec1. For all of figure 4, images of virus-infected cells were aggregated from 2 independent experiments, each with at least 10 technical replicates per treatment.

Logit scores, which quantify the probability that a given cell falls into each class, from dying, virus-infected cells primarily overlap with prototypical apoptotic and necroptotic cells, with a much smaller fraction falling into the pyroptosis cluster (Fig 4C). These data also recapitulate the time-dependent shift of cells from the necroptotic to apoptotic fate. Over the course of 48 hours, roughly 38%, 53%, and 8% of cells are classified as apoptotic, necroptotic, or pyroptotic, respectively (Fig 4E). These data demonstrate that HSV-1 infected cells execute variable and dynamic modes of PCD.

It has been proposed that crosstalk between the different PCD pathways allows dying cells to flexibly switch to a different pathway if one is blocked (Snyder and Oberst, 2021). To test the flexibility of infected cells in switching between death pathways, we infected cells with HSV-1 and treated them with either a caspase inhibitor (ZVad) or a RIP1K inhibitor (Nec1). Caspase inhibition triggered significantly faster, yet reduced overall cell death compared to cells treated with the vehicle control (Fig 4D). Conversely, inhibiting RIP delayed the onset of cell death compared to the vehicle control, although a similar overall fraction of cells eventually died by 48 hours post-infection. These results show that, contrary to the idea of full redundancy between these pathways, caspase- and RIPK-dependent forms of PCD are not able to substitute for each other during infection, indicating that these death programs are not fully compensatory.

We next used our classifier to test whether the application of caspase or RIPK inhibitors alters the proportion of cells undergoing each type of cell death. Compared to the control, caspase inhibition significantly reduced the fraction of cells classified as apoptotic and increased the proportion classified as necroptotic (Fig 4E). Conversely, although RIP inhibition delays cell death, it does not affect the overall frequency of cells classified as necroptotic throughout the experiment (Fig 4D-E).

Finally, we used our cell death classifier to evaluate the kinetics of the different PCD pathways over time and with pharmacologic inhibition of either RIP or caspases. In control cells, necroptosis strongly predominates early during infection, whereas the proportion of apoptotic cells increases over time (Fig 4A, C F-G). While caspase inhibition potently favors necroptosis over apoptosis in the cells that died, the overall reduction in death implies that cells that were destined to apoptosis simply survived, rather than switched to necroptosis. Adding a RIP inhibitor delayed the onset of both necroptosis and apoptosis, but did not significantly alter the fraction of cells dying by different modes. The modest effect of RIP inhibition on apoptosis is consistent with reports that RIP can direct a caspase-independent form of apoptosis specifically during viral infection (Nogusa *et al*., 2016). Neither caspase nor RIP inhibition altered the proportion of cells that undergo pyroptosis (Fig 4H).

Overall, these data show that viral infection triggers diverse PCD pathways within a uniform population of cells. Necroptosis is the dominant mode of cell death early in the infection, but apoptosis becomes more prevalent as the infection progresses. While inhibiting caspases shifts the proportion of death modes from apoptosis to necroptosis, that shift can be explained by a fractional killing effect, where apoptosis-destined cells survive. Inhibiting RIP simply delays cell death without changing the type of PCD. Additionally, neither treatment affected the proportion of cells undergoing pyroptosis. These findings suggest that, although the population as a whole exhibits plasticity in the activation of different PCD pathways, individual cells show limited flexibility in switching between modes. Therefore, distinct PCD mechanisms provide overall adaptability during infection, but single cells are constrained in their ability to alter their death fate.

## Discussion

Molecular measurements of signaling pathways and gene expression are the workhorse of biological research, but can fall short in determining cell fate. This is especially true when multiple cell signaling pathways are activated in parallel, but only one is fully executed. We found that programmed cell death (PCD) during viral infection is a prime example of this complexity, as many infected cells show activation of both apoptotic and necroptotic signaling pathways (Fig 1). Historically, microscopy-based morphology assessments could be used to distinguish execution of these two different PCD types, but were limited by lower throughput and reliance on human expertise. Recent advances in computer vision now offer an opportunity to integrate morphological data back into biological research in a systematic and scalable way (Moen *et al*., 2019).

To build a robust morphology-based classifier, it was crucial to create accurate training data that captured heterogeneity without introducing mislabeled samples. We treated cells with drugs that selectively induce apoptosis, necroptosis, or pyroptosis, then used automated image analysis and confident learning to prune incorrect labels from the dataset (Northcutt *et al*., 2019) (Figs 2-3). By training a residual neural network (ResNet) on this refined data, we were able to accurately classify cells based on their mode of death (Fig 3) (He *et al*., 2015). This approach builds on past work that utilized feature extraction, as opposed to pixel-level classification - or holographic microscopy to distinguish different death modes (Barker *et al*., 2020; Vicar *et al*., 2020; Verduijn *et al*., 2021). Our results showed that HSV-1-infected fibroblasts primarily died via necroptosis early on, with apoptosis becoming more prevalent later in infection (Fig 4). Importantly, inhibiting caspases did not cause cells to switch to necroptosis, and vice-versa, challenging the notion of functional redundancy in death pathways during infection.

This approach overcomes the limitations of manually annotated datasets, which tend to be morphologically uniform and less representative of real data variability. By combining a pharmacological strategy with computational dataset pruning, we generated a more accurate classifier. Our findings suggest that cells are predisposed to specific death modes and have limited ability to switch when one pathway is blocked. This “bet-hedging” strategy might help the population resist viral spread, as some infected cells will die regardless of which pathway the virus targets. Our experimental and computational framework could be extended to study other processes involving fate-associated morphologies, such as mitosis, differentiation, and others. In summary, deep learning-based image classification improves the accuracy and scalability of cell death assignment from microscopy data. We expect this approach to enhance and complement existing molecular assays that probe for biochemical evidence of distinct PCD fates.

## Methods

### Cells and Culture Conditions

NIH 3T3 and Vero cells were purchased from the American Type Culture Collection (ATCC) (#CRL-1658 and #CCL-81), and grown at 37°C and 5% CO_2_ in a humidified incubator. Cells were grown in MEM, supplemented with 10% fetal bovine serum, 1% penicillin/streptomycin, and 2mM L-glutamine.

### Cytotoxic prototype treatments

To induce apoptosis, cells were treated with a combination of 10 ng/ml mouse TNFα (Peprotech), 30μM Necrostatin-1 (Sigma-Aldrich); and 3 μg/mL Cycloheximide (Sigma-Aldrich). To induce necroptosis, cells were treated with a combination of 10 ng/ml mouse TNFα, 30μM Z-VAD(Ome)-FMK (Cayman Chemical), and 3μg/mL Cycloheximide. To induce pyroptosis, cells were treated with a combination of 250ng/ml K12 E. coli-derived LPS (Invivogen) and 10μM Nigericin (Invivogen).

### Viruses and viral propagation

HSV-1 strain KOS bearing fluorescent fusion proteins (Etienne *et al*., 2017) of the small capsid protein VP26 to tdTomato, mCherry, or mCerulean were provided by Dr. Prashant Desai (Johns Hopkins University). To propagate viruses, 4 × 10^6^ Vero cells were seeded in T150 cm^2^ tissue culture flasks, then infected with HSV-1 strains at an MOI of 0.05. To infect, medium was removed and a 1.5 mL suspension of virus in PBS was added. After rocking briefly, the flask was transferred to 37°C and rocked every 10 min. After 1h, 45 mL of growth medium was added and incubated for 2 days at 37°C. After 2 days, cells were detached by shaking/tapping, and then pelleted for 20 minutes at 4°C. The supernatant was transferred to a conical tube and kept on ice. The remaining cell pellet was resuspended in 1 mL of growth medium, and cells were lysed by repeatedly pushing through a 1.5 mL syringe. 8 mL of growth medium was added to the lysate, and clarified by spinning for 10 minutes at 4°C. The two supernatant sources were then combined and centrifuged in a Beckman Coulter Optima L-100XP Ultracentrifuge at 40,000 g for 30 minutes at 4°C. After centrifugation, the supernatant was removed, and the remaining pellet was resuspended in MEM supplemented with 10% FBS and 10% glycerol to a final volume of 1.8 mL. The virus suspension was then homogenized using a 1.5 mL syringe and stored at -80°C. Viral titers were quantified by standard plaque assays performed in Vero cells.

### Immunofluorescence Staining

For immunofluorescence, 3.5 × 10^4^ cells were cultured in Nunc Chambered Coverglass 8 well Lab-Tek™ II plates. Cells were fixed with 1.6% paraformaldehyde for 10 min on ice, then washed with PBS and permeabilized with ice-cold 90% methanol for at least 1 hour. For antibodies directed against pRIP or pMLKL, cells were fixed only with ice-cold 90% methanol (Samson *et al*., 2021). Fixed and permeabilized cells were then washed 3x with PBS, and blocked for 1h at room temperature. Blocking buffer consists of PBS, 0.3% Triton X-100, and 5% normal human serum (cat: 009-000-121, Jackson Immunoresearch). Human serum is essential to block non-specific binding of primary antibodies to HSV-1 infected cells. Cells were stained with antibodies for 30m-1h at room temperature or up to overnight at 4°C. Where, indicated, cells were then washed 3x with blocking buffer, then incubated with secondary antibodies and counterstained with DAPI for 1h at room temperature, light protected. Cells were then washed 3x with blocking buffer and an additional 2x with PBS before imaging.

#### Primary antibodies

rabbit anti-pMLKL (clone D6E3G, Cell Signaling Technologies), rabbit anti-pRIP (clone E9K2A, Cell Signaling Technologies), rabbit anti-Cleaved Caspase 3 (clone 9661, Cell Signaling Technologies), Rabbit anti-Cleaved Caspase 8 (clone D5B2, Cell Signaling Technologies), mouse anti-ICP4 (clone 10F1, Abcam), rabbit IgG isotype control antibody (clone DA1E, Cell Signaling Technologies)

#### Secondary antibodies

donkey anti-goat F(ab’)_2_-Alexa647 (cat: 705-606-147, Jackson Immunoresearch), goat anti-rabbit F(ab’)_2_-Alexa488 (cat: 111-546-003, Jackson Immunoresearch), goat anti-mouse F(ab’)_2_-Alexa647 (cat: 115-606-003, Jackson Immunoresearch)

### Fixed Cell Microscopy

Images were collected on a confocal Nikon Eclipse Ti2E-inverted microscope equipped with a Nikon CFI Plan Apo λ 20×, numerical aperture (NA) 0.75 objective lens. Images were acquired with a CrestOptics X-Light V3 light engine and a Photometrics Kinetix camera. All imaging was accomplished using custom automated software written using MATLAB and Micro-Manager.

### ATP Release Assay

The RealTime-Glo Extracellular ATP Assay Reagent was mixed with 10 mL of culture medium (RPMI 1640, supplemented with 10% fetal bovine serum and 1% penicillin/streptomycin (volume:volume) yielding a 4X reagent. 6.00 × 10_4_ cells were seeded in culture medium in Greiner CELLSTAR 96 well plates and allowed to adhere for 24 hours. Cells were then treated with compounds, and 50µl of 4X RealTime-Glo Extracellular ATP Assay Reagent was added to each well. Cells were then imaged using a BioTek Synergy HTX plate reader.

### Time-lapse Imaging

3.5 × 10^4^ cells were seeded in imaging media (MEM - no phenol red / no glutamine, 10% FBS, 10U/ml Penicillin and 10 μg/ml Streptomycin) in Nunc Lab-Tek II Chambered Coverglass 8 well or 2.5 × 10^4^ cells were seeded in 96 well glass bottom plate with high performance #1.5 cover glass (CellVis). Cells were labeled with 20 ng/ml Hoechst 33342 (Thermo Fisher) and death was tracked by incorporation of 1:10,000 Sytox Green (Thermo Fisher). Images (20×) were collected on a Nikon Eclipse Ti2-E inverted microscope equipped with a Nikon CFI Plan Apo Lambda 20×, numerical aperture (NA) 0.75 objective lens. Images were acquired with a Lumencor SOLA SE V-nIR light engine and Hamamatsu ImagEM EM-CCD camera. NIS Elements software was used for image acquisition.

### Timelapse Image Analysis

Automated image processing and analysis was performed using custom code written in Python. Software is available from the GitHub repository: https://github.com/oylab/oyLabImaging. Briefly, phase-correlation based jitter and drift correction was applied to timelapse image stacks. Single cells were identified and segmented using the pretrained segmentation model stardist (version 0.8.3, (Schmidt *et al*., 2018)), and single cell features were saved. Single cells were then tracked using our custom made tracking software based on the python package **lap05** (Jonker and Volgenant, 1987). Cell death was identified as a sustained (>4 consecutive frames) increase of either the intensity of the nuclear marker Hoechst 33342, indicative of nuclear condensation, or Sytox Green, indicative of membrane rupture, above a user-selected threshold. 128×128 pixel DIC images of individual cells taken from the frame immediately prior to death were saved for deep learning model training.

### ResNet models

Model training and deployment were performed using custom Python code. We implemented transfer learning by using the pretrained ResNet50 model as our base (https://download.pytorch.org/models/resnet50-0676ba61.pth). The model was adapted to accept single 128×128 grayscale inputs and trained on images of imminently dying cells exposed to prototype-inducing drugs. Training was done using stochastic gradient descent (SGD) with Focal Loss (gamma=2) (Lin *et al*., 2017). We trained 10 independent models on an identical dataset with a 70/15/15 split for training, validation, and testing, with random shuffling. Each model was trained for 150 epochs, using a minibatch size of 32 for the training set and 400 for the validation and testing sets. The training set was class-balanced.

After initial training, we pruned incorrectly labeled samples by removing any sample that was not classified as its ground truth label by at least 80% of the models (using softmax probability estimates) across the initial 10 models. The pruned dataset was then used to train a new model using standard Cross Entropy as the loss function and SGD over 500 training epochs. Deployment of the ResNet models was also done with custom Python code, based on the PyTorch distribution. The software is available on GitHub: https://github.com/oylab/oyDL.

### Statistical Analysis and Data Presentation

To visualize our classifier outputs, we performed dimensional reduction using Uniform Manifold Approximation and Projection (UMAP). We generated UMAP representations of the final classifier features by extracting activations from the last fully connected layer, which contains 2048 features measured over the entire pruned dataset. A 2D UMAP representation was fit using 100 neighbors and a minimum Euclidean distance of 0.1. Features extracted from virus infected cells were projected onto the same prototype projection, and were not fit independently.

All relevant data are shown as mean ± standard error of the mean (s.e.m.) or standard deviation (s.d.). For microscopy images, identical brightness and contrast settings are used in side-by-side images.

## Author Contributions

Conceptualization, J.O.-Y., and A.O.-Y.. Investigation, E.C., A.O.-Y. and J.O-Y. Software, A.O.-Y. Writing, J.O.-Y., A.O.-Y., and E.C. Resources, J.O.-Y.

## Acknowledgements

We thank the Core for Imaging Technology and Education (CITE, HMS) for providing access to microscopes and consultation on imaging conditions, Dr. Talley Lambert (CITE, HMS) for helping with image analysis software development, Dr. Prashant Desai (Johns Hopkins University) for sharing viruses and protocols for their propagation, Drs. Tim Mitchison (HMS), Carlos Lopez (Altos Labs), and all members of the Oyler-Yaniv lab for helpful conversations, and Karla Flores for lab support.

## Figure Legends

**Supplementary Figure 1.**
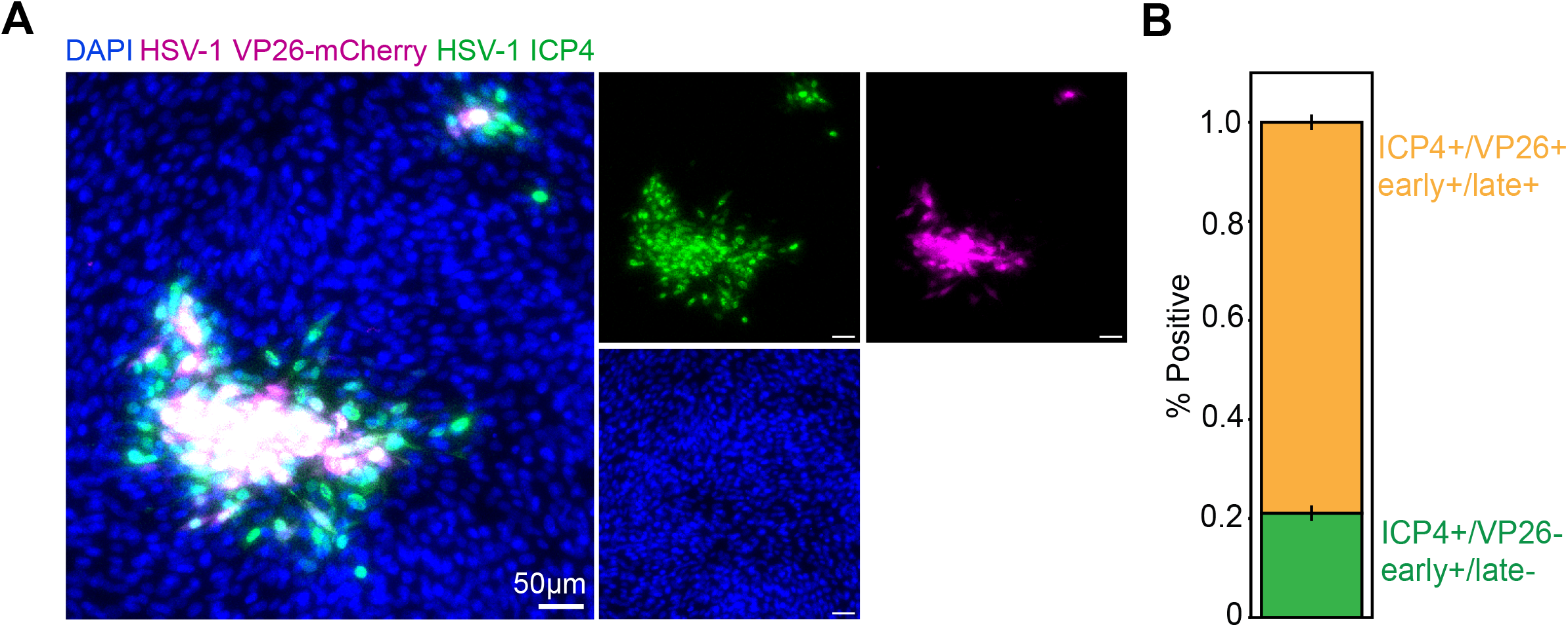
A small proportion of early-infected cells are negative for viral VP26. **A**. Representative immunofluorescence images of mouse fibroblasts infected with HSV-1 strain VP26-mCherry and stained for the viral immediate-early protein ICP4. **B**. Quantification of the proportion of cells that are either ICP4-positive and VP26-negative, or double positive for both proteins. Error bars show the standard deviation for three biological replicates.

**Supplementary Figure 2.**
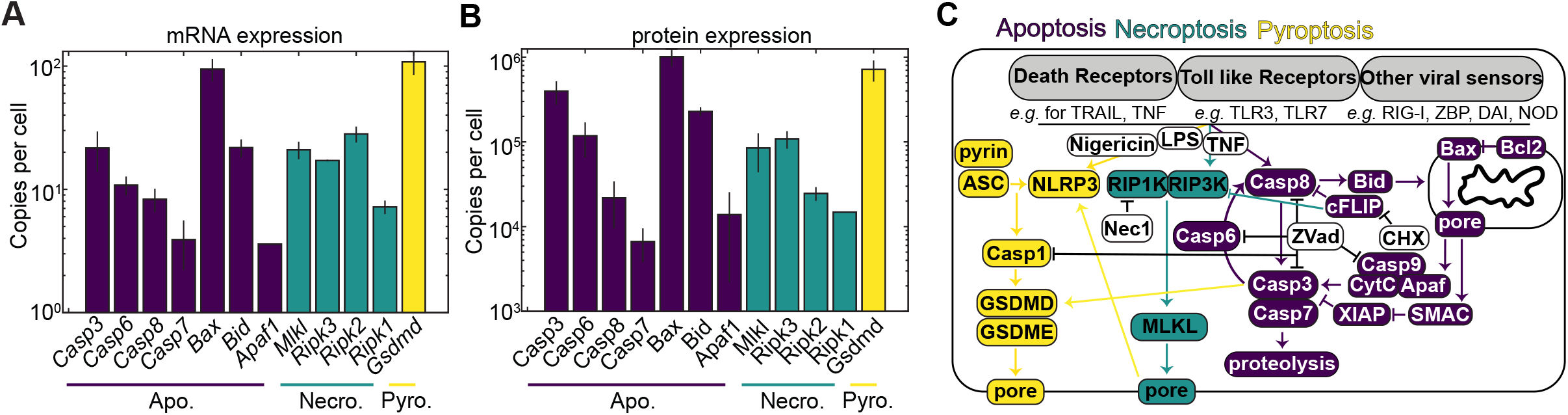
Key components of each of the three major PCD signaling pathways. **A**-**B**. mRNA (A) and protein (B) expression values for core components of each PCD signaling pathway for 3T3 fibroblasts from (Schwanhäusser *et al*., 2011). **C**. Cartoon diagram of the signaling cascades that drive apoptosis, necroptosis, and pyroptosis. In white are the pharmacologic components that are used to activate each of the PCD types in our experiments. Error bars in A and B represent the standard deviation of two replicate experiments.

**Supplementary Figure 3.**
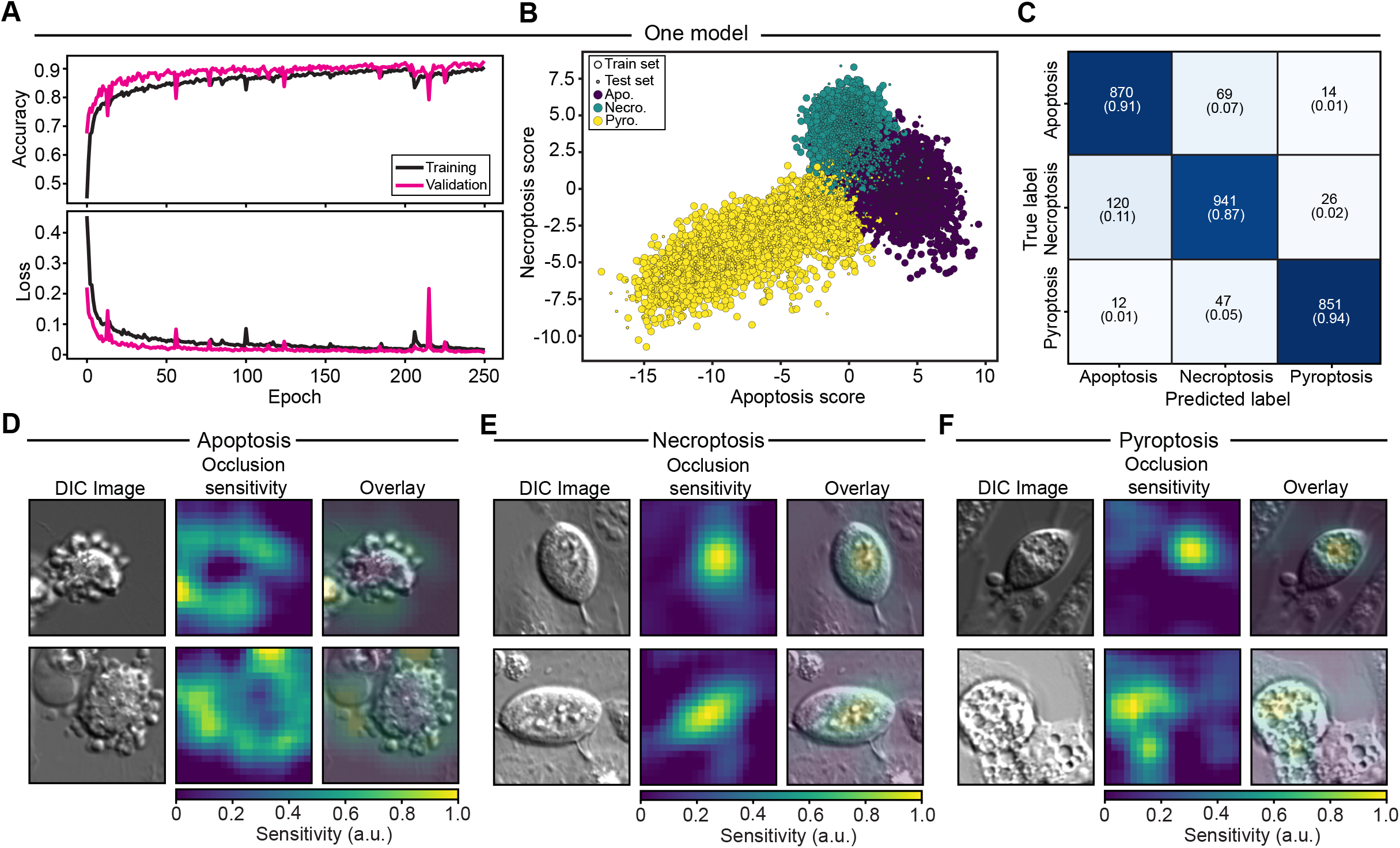
Pruning the labeled data using consensus models improves model training. **A**. Performance of the classifier during training before employing any consensus-based pruning. **B**. Logit scores of the train and test data in absence of pruning. **C**. Confusion matrix demonstrating model performance in absence of pruning. Numbers represent the number of cells classified (top) and fraction called correctly (bottom). **D-F**. Occlusion sensitivity for additional representative cells from each death prototype class.

